# Integrating Fossil Flowers into the Angiosperm Phylogeny using a Total Evidence Approach

**DOI:** 10.1101/2022.02.17.480913

**Authors:** Andrea M. López-Martínez, Jürg Schönenberger, Maria von Balthazar, César A. González-Martínez, Santiago Ramírez-Barahona, Hervé Sauquet, Susana Magallón

**Affiliations:** Posgrado en Ciencias Biológicas, Instituto de Biología, Universidad Nacional Autónoma de México, 3er Circuito de Ciudad Universitaria, Del. Coyoacán, Ciudad de México 04510, Mexico; Departamento de Botánica, Instituto de Biología, Universidad Nacional Autónoma de México, 3er Circuito de Ciudad Universitaria, Del. Coyoacán, Ciudad de México 04510, Mexico; Department of Botany and Biodiversity Research, University of Vienna, Rennweg 14, Vienna A-1030, Austria; National Herbarium of New South Wales (NSW), Royal Botanic Gardens and Domain Trust, Sydney, NSW 2000, Australia; Evolution and Ecology Research Centre, School of Biological, Earth and Environmental Sciences, University of New South Wales, Sydney, Australia

**Keywords:** Angiosperms, Fossil Flowers, Phylogenetic Uncertainty, RoguePlots, Total Evidence

## Abstract

Fossil flowers are essential to infer past angiosperm evolutionary processes. The assignment of fossil flowers to extant clades has traditionally relied on morphological similarity and on apomorphies shared with extant taxa. The use of explicit phylogenetic analyses to establish their affinity has so far remained limited. In this study, we built a comprehensive framework to investigate the phylogenetic placement of 24 exceptionally preserved fossil flowers. For this, we assembled a new species-level dataset of 30 floral traits for 1201 extant species that were sampled to represent the stem and crown nodes of all angiosperm families. We explored multiple analytical approaches to integrate the fossils into the phylogeny, including different phylogenetic estimation methods, topological-constrained analyses, and a total evidence approach combining molecular and morphological data of extant and fossil species. Our results were widely consistent across approaches, with minor differences in the support of fossils at different phylogenetic positions. The placement of some fossils is in agreement with previously suggested relationships, but for others, a new placement is indicated. We also identified fossils that are well constrained within particular extant families, whereas others showed high phylogenetic uncertainty. Finally, we present recommendations for future total evidence analyses, regarding the selection of fossils and appropriate methodologies, and provide some perspectives on how to integrate fossils into the investigation of divergence times and the temporal evolution of morphological traits.

## Main

The information provided by the fossil record is essential to infer diverse evolutionary processes in deep time (Marshall 2017; Benson et al. 2021). It is needed to determine the origin of major living groups (e.g., O’Leary et al. 2013; Misof et al. 2014; Magallón et al. 2015), to understand diversification dynamics (Silvestro et al. 2015; Mitchell et al. 2019; Lloyd and Slater 2021), to reconstruct ancestral states (Slater et al. 2012; Betancur-R et al. 2015; Larson-Johnson 2016), or to infer the biogeographical history of lineages (Meseguer et al. 2015; Zhang et al. 2021). The recent development of novel methodological approaches has made possible a more integrative use of the fossil information, such as by the incorporation of fossils into the same diversification process as extant species through the Fossilized Birth-Death model (Heath et al. 2014); or by using the fossil record to estimate changes in origination and extinction rates using Bayesian approaches (Silvestro et al. 2014; Mitchell et al. 2019). However, these approaches share a requirement of *a priori* specifications about the taxonomic assignment of the fossils, which in turn are based on morphological information that, in the best-case scenario, are also supported by phylogenetic analyses. The tip dating approach, in which fossils appear as extinct tips within a phylogenetic tree, uses fossil ages and total evidence matrices to simultaneously infer the placement of fossils and calibrate the tree to estimate divergence times (Ronquist et al. 2012; Zhang et al. 2016; Gavryushkina et al. 2017). Tip dating is a promising approach but it requires the assemblage of total evidence matrices that integrate molecular data of extant species, with morphological data scored for both extant and fossil taxa. The assembly of such matrices is particularly challenging in broad-scale analyses because of the difficulty of generating robust and comprehensive data for a large sample of species. The phylogenetic investigation of fossils using total evidence data has so far remained limited in part due to the lack of morphological matrices for a broad sample of extant and fossil taxa, possible methodological artifacts linked with large amounts of missing data for fossils (Manos et al. 2007), and by the use of simplified models of morphological character evolution that result in distorted topologies and branch lengths (Ronquist et al. 2016; Wright et al. 2016).

Fossil flowers are an invaluable source of information to comprehend floral morphological diversification and the temporal emergence of novel structures across angiosperms. These fossilized flowers also provide insights into the diverse evolutionary mechanisms in angiosperms and their interactions with pollination and dispersal vectors (Friis et al. 2010, 2011). Although flowers are preserved through different processes such as permineralization, impressions/compressions, and amber, it is the charcoalified flowers that have been most extensively studied (Friis et al. 2006, 2011). Charcoalified flowers are most commonly represented in Cretaceous sediments, possibly due to increases in paleo-fire regimes during that period (Bond and Scott 2010). This type of fossils have been described mainly from localities in the Eastern United States (Drinnan et al. 1990, 1991; Crepet and Nixon 1998; Gandolfo et al. 1998; von Balthazar et al. 2007; Crepet et al. 2018; Friis et al. 2020), Portugal (Friis et al. 2001, 2019), Sweden (Friis and Skarby 1981; Schönenberger and Friis 2001; Friis and Pedersen 2012), the Czech Republic (Heřmanová et al. 2021, 2022), and Japan (Takahashi et al. 2001, 2008). The preservation of charcoalified flowers is often exquisite, allowing for the detailed study of morphological and anatomical floral traits. Thanks to their rigidity and resistance against compression, many of these charcoalified specimens are preserved in their original 3D-shape, frequently preserving internal and external structures in great detail (Schönenberger 2005). Revolutionary visualization techniques, such as synchrotron radiation x-ray, have improved the quality and quantity of morphological information that can be obtained from the fossil record (Friis et al. 2014). Therefore, internal morphological details of charcoalified flowers may be obtained at anatomical or cellular resolution, equivalent to living species.

The systematic assignment of fossil flowers has so far mainly relied on extensive morphological investigations and on the identification of apomorphies shared with living taxa (e.g., Drinnan et al. 1990; Magallón et al. 2001; von Balthazar et al. 2005). Studies that identify or corroborate systematic hypotheses of fossil flowers using phylogenetic methods are scarce (e.g., Keller et al. 1996; Gandolfo et al. 2002; Magallón 2007; Doyle and Endress 2010, 2014; Lee et al. 2013; Martínez et al. 2016; Schönenberger et al. 2020). Most of these analyses investigate the position of individual fossil flowers by focusing on a specific subclade of angiosperms (i.e., an order, a family, or a genus), which has been defined *a priori* based on comparative morphology and taxonomic expertise (e.g., Hermsen et al. 2003; Mendes et al. 2014; Martínez et al. 2016). Consequently, these analyses exclude the possibility that the fossil under study falls outside of the focal group (widely discussed by Schönenberger et al. 2020). Relatively few studies have applied a more broad-scale approach to cover multiple major angiosperm subclades, including Doyle and Endress (2010, 2014), and the broadest one, published recently by Schönenberger et al. (2020). In these studies, the fossils are integrated into a molecular backbone phylogeny of extant species based on parsimony, without exploring alternative methods or optimization criteria. In addition, the phylogenetic placement of fossils was assessed one fossil at the time, thereby removing the potential benefit of the interaction between multiple fossils and extant species.

Here, we integrate molecular and morphological data in a comprehensive framework to jointly analyze the phylogenetic placement of 24 well-preserved fossils flowers across the entire angiosperm phylogeny. For this, we assembled the largest dataset of floral traits ever published, and considerably expanding on previous datasets (Sauquet et al. 2017; Schönenberger et al. 2020). This dataset includes extant representatives of all currently recognized angiosperm families *sensu* the Angiosperm Phylogeny Group IV (APG IV, 2016) and mirrors the sampling of species in the molecular dataset of Ramírez-Barahona et al. (2020). We use diverse phylogenetic approaches to assess the placement of fossils and subsequently visualize their phylogenetic uncertainty. We implement previous approaches based on the application of topological constraints among extant species and the analysis of each fossil independently, as well as new unconstrained analyses based on total evidence matrices for the phylogenetic estimation of multiple fossil and extant species simultaneously. Finally, we provide recommendations for the investigation on the phylogenetic placement of fossils based on analyses of combined molecular and morphological data.

## MATERIALS AND METHODS

### Extant and Fossil Taxon Sampling

We selected a total of 24 exceptionally well-preserved fossils flowers with the aim of having a broad representation of fossils across all major lineages of angiosperms (according to the systematic positions established in earlier studies; Table 1 and Supplementary Information Data 1). Most of the fossils selected for this study are three-dimensionally preserved charcoalified flowers (e.g., Crane et al. 1989; Magallón-Puebla et al. 1997; von Balthazar et al. 2008). However, we also included permineralized fossils (e.g., Smith and Stockey 2007) and impressions/compressions (e.g., Manchester 1992). The set of fossils included the ten fossil flowers that were analyzed in the study by Schönenberger et al. (2020) with the aim of comparing their phylogenetic placements using different sampling and methods. We used the molecular dataset of extant species from Ramírez-Barahona et al. (2020), which consisted of seven molecular markers: four protein-coding plastid genes (*rbcL*, *atpB*, *matK*, *ndhF*); and three nuclear loci (*18S*, *26S*, and *5.8S* nrDNA). The original study by Ramírez-Barahona et al. (2020) encompasssed 1209 taxa, including a combination of explicit species and chimeric taxa (whereby different genes were sampled from multiple species within the same genus to maximise data coverage). Here we used the same approach as Sauquet et al. (2017) to match each of these taxa in the molecular dataset to an explicit species in our morphological dataset. In this process, eight taxa could not be matched or became redundant, leading to a molecular dataset and tree of 1201 extant species representing the stem and crown nodes of all angiosperm families *sensu* APG IV (2016) and the Angiosperm Phylogeny Website (http://www.mobot.org/MOBOT/research/APweb/; Stevens, 2001).

**Table 1.**
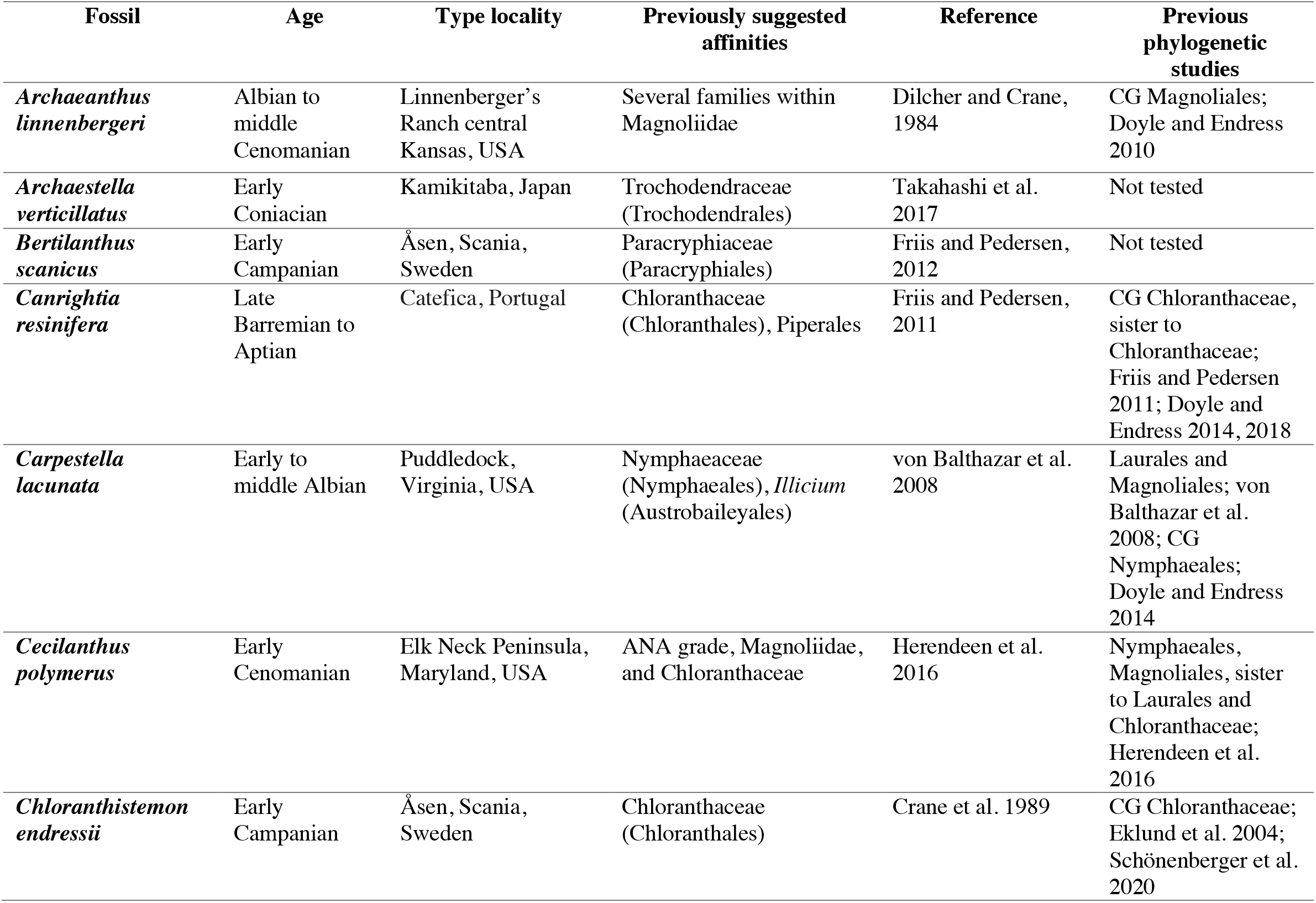
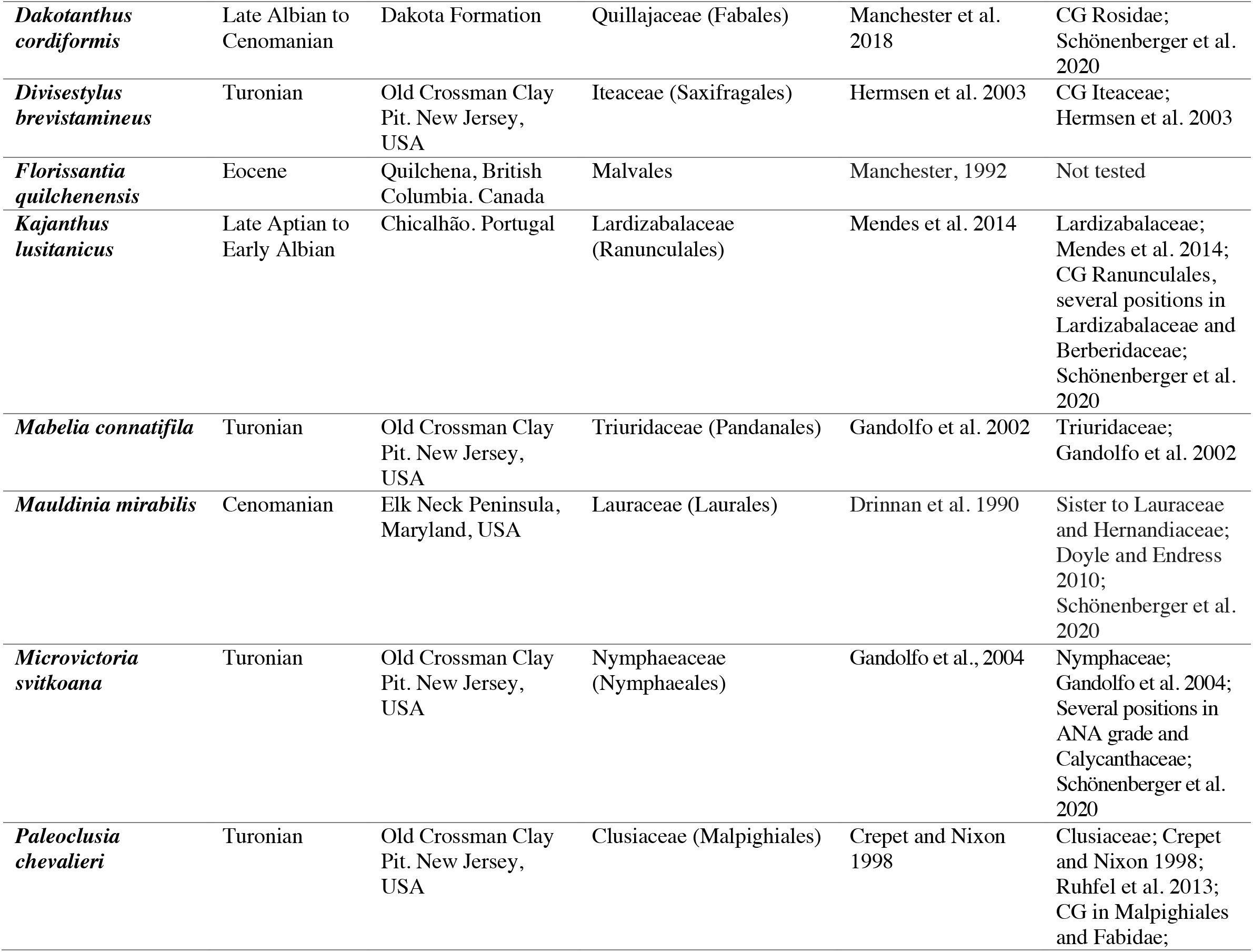
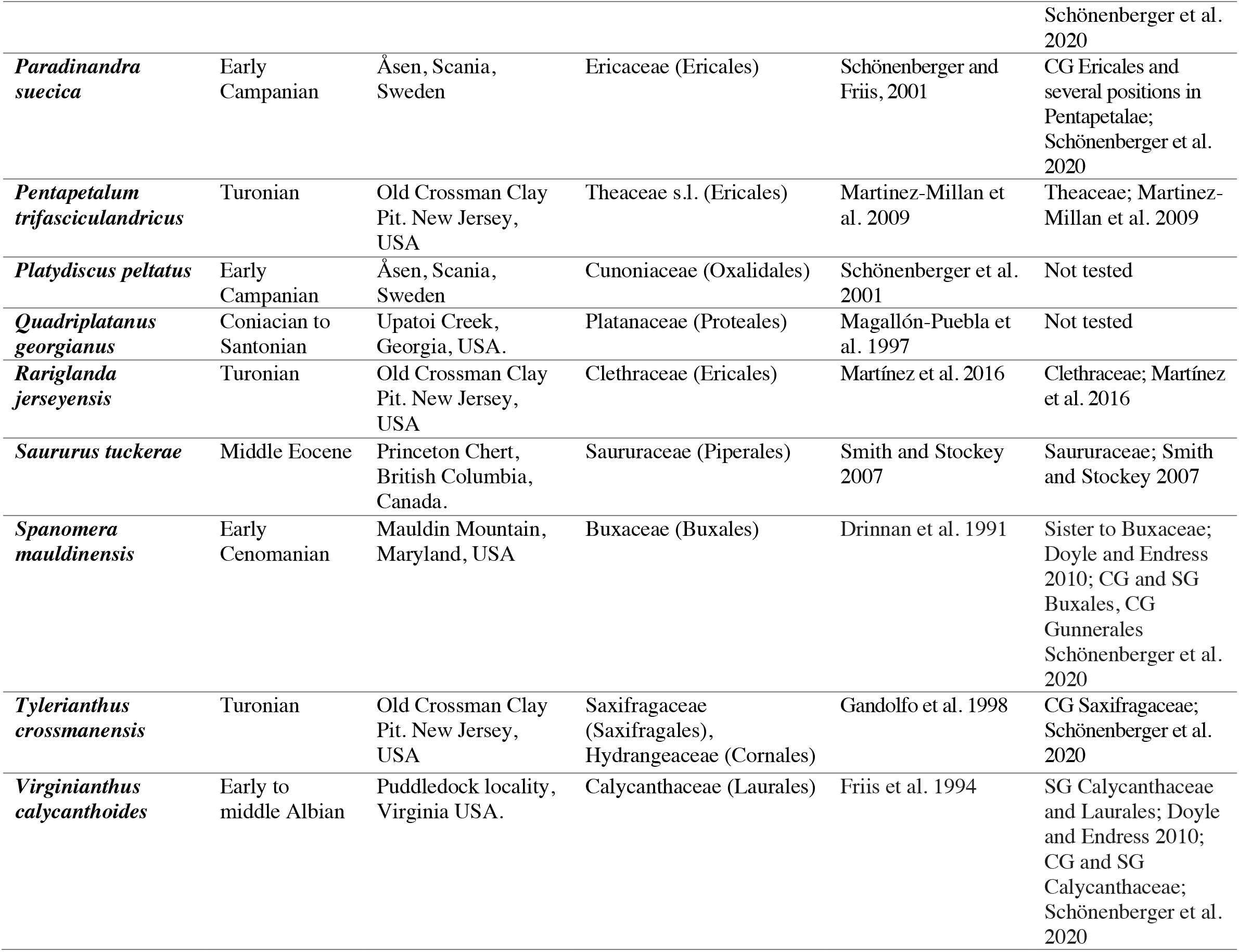
Main information about age, locality, previously suggested affinities of the 24 fossils included in this study. CG, Crown group; SG, Stem group.

### Morphological Data

The morphological dataset consisted of 30 floral traits recorded for 1201 extant and 24 fossil species in the PROTEUS database (Sauquet 2019). This is the same set of floral characters defined and used by Schönenberger et al. (2020), including structural traits describing the sex of flowers (1 character), the perianth (8 characters), the androecium (11 characters), the gynoecium (8 characters), and pollen (2 characters). For details on our scoring philosophy and definition of characters see published appendices in Sauquet et al. (2017) and Schönenberger et al. (2020). The morphological traits were scored based on an extensive search of data in a wide range of published literature, including original papers, text books, and floras. In total, 567 extant and 10 fossil species are shared between this new dataset (of 1201 extant and 24 fossil species) and previously published datasets (of 792 extant and 10 fossil species; Sauquet et al. 2017; Schönenberger et al. 2020). We curated and filled-in gaps in these published datasets and hence, out of 13,812 total data records for these overlapping species, 13,730 were previously published, 82 are new, 169 were updated, and 52 were deleted. The majority of the remaining 634 extant and 14 fossil species were scored here for the first time, corresponding to a total of 12,785 new data records. The complete list of 26,597 total data records, along with explicit references (in total 2020 distinct sources) used to score each species is provided as Supplementary Information Data 2. As in previous publications based on the PROTEUS database, the data were recorded as primary continuous or discrete characters, and then transformed into a matrix of secondary, discrete characters for the purpose of analysis (see Sauquet et al. 2017, Schönenberger et al. 2020). The final morphological matrix (Supplementary Information Data 3) consists of 36,750 cells with a proportion of missing data of approximately 32% and 23% for extant and fossil species, respectively.

### Constrained Analyses

To estimate the phylogenetic placement of fossils, we conducted multiple independent phylogenetic analyses (summarized in Table 2). The first type of analyses incorporated a molecular backbone tree to fully constrain the topology of extant species. For this purpose, we selected one of the maximum credibility time-trees obtained by Ramírez-Barahona et al. (2020) (Supplementary Information Data 4). This tree was reconstructed in BEAST based on the ‘relaxed calibration strategy’ described in (Ramírez-Barahona et al. 2020), which included one prior constraint on the crown age of angiosperms and 238 fossil-based minimum age constraints. First, we estimated the phylogenetic position of each fossil independently following the *phyloscan* (here *CMP-1-Morph*) approach as implemented by Schönenberger et al. (2020). For these analyses, a parsimony score was calculated for each fossil (independently of other fossils) for every possible placement in the tree (all terminal and internal branches, a total of 1201+1199), based on the morphological matrix scored for the fossil of interest and all extant representatives. Each of the 24 *CMP-1-Morph* analyses were conducted with the R script published by (Schönenberger et al. 2020) using the package *phangorn* v.2.5.2 (Schliep 2011). Second, we conducted an analysis for each fossil as described for *CMP-1-Morph* but with a maximum likelihood (ML) optimization criterion (here *CML-1-Morph*). All ML analyses were performed with RAxMLv.8.0 (Stamatakis 2014), implementing the constrained topology among extant species with the option -g, and the ASC_MULTICAT model with an ascertainment bias correction=lewis (Lewis 2001) for morphological data. A total of 1000 replicates of non-parametric bootstrap were obtained. Third, we ran a single ML analysis incorporating the 24 fossil flowers simultaneously, using the morphological matrix for extant and fossil species (*CML-24-Morph*). This analysis was conducted with the same parameters used for each of the *CML-1-Morph* analyses. Fourth, we used ML to estimate the placement of all fossils simultaneously, using the total evidence matrix consisting of molecular data for extant species and morphological data for extant and fossil species (*CML-24-TE*). As the topology of the tree remained constrained, this analysis was aimed at investigating the effects of molecular branch lengths as well as missing molecular data for fossils on their phylogenetic placement. ML phylogenetic estimation was conducted with RAxML v.8. (Stamatakis 2014), using the ASC_MULTICAT model with an ascertainment bias correction=lewis (Lewis 2001) for morphological data, and the GTR model for the following molecular partitions: *rbcL* 1^st^ and 2^nd^ positions; *rbcL* 3^rd^ position; *atpB* 1^st^ and 2^nd^ positions; *atpB* 3^rd^ position; *matK; ndhF*; and *nuclear*, as specified by Ramírez-Barahona et al. (2020). We ran 1000 non-parametric bootstrap replicates. All phylogenetic analyses were performed in the CIPRES science gateway (Miller et al. 2010).

**Table 2.** List of phylogenetic analyses conducted in this study.

| Constraint | Optimization Criterion | Taxa subjected to phylogenetic estimation | Data | Analysis code |
| --- | --- | --- | --- | --- |
| **Backbone** (relationships among extant taxa derived from molecular data) | parsimony | 1 fossil (one analysis for each of the 24 fossils) | morphological | *CMP-1-Morph* |
|  | ML | 1 fossil (one analysis for each of the 24 fossils) | morphological | *CML-1-Morph* |
|  | ML | 24 fossils simultaneously | morphological | *CML-24-Morph* |
|  | ML | 24 fossils simultaneously | total evidence | *CML-24-TE* |
| **Unconstrained** | ML | 24 fossils & 1201 extant species simultaneously | total evidence | *UML-24-TE* |
|  | BI | 24 fossils & 1201 extant species simultaneously | total evidence | *UBI-24-TE* |

### Unconstrained Analyses

In the second set of analyses, no backbone tree was used to constrain the relationships among extant species, thus allowing direct phylogenetic estimation among extant and fossil species. We estimated phylogenetic relationships among extant and fossil species using the total evidence matrix described above, with ML and Bayesian Inference (BI). The unconstrained ML analysis (*UML-24-TE*) was conducted with RAxML v.8. (Stamatakis 2014), implementing the same set of partitions as in *CML-24-TE*. In the unconstrained BI analyses (*UBI-24-TE*), we initially did not fix the relationships among extant taxa, but due to problems in MCMC convergence, and following Ramírez-Barahona et al. (2020), we constrained some clades to be monophyletic (Supplementary Information Data 5). BI was performed using MrBayes v3.2.7 (Ronquist et al. 2012b) implementing the Mk Markov Model (Lewis 2001) for categorical data and the same partitions as those used in the ML analyses for molecular data. Two independent runs of four chains with a temperature of 0.2 were run for 30 million MCMC generations, sampling one tree each 5000 steps. MCMC convergence was visualized in Tracer v 1.7. 2 (Rambaut et al. 2018), checking that ESS values were above 200 for all parameters.

### Visualization of Phylogenetic Uncertainty

We constructed RoguePlots (Klopfstein and Spasojevic 2019) as a graphical aid to visualize phylogenetic uncertainty in the placement of fossils in ML and BI phylogenetic analyses. This function summarizes all possible attachments of fossils to branches in the tree, each supported by its posterior distribution or bootstrap replicates. Instead of relying on a single summary position for a fossil, this method shows all sampled phylogenetic placements of a given fossil across the tree weighted by their support value. The RoguePlots graphs were constructed using the package *rogue.plot* (Klopfstein and Spasojevic 2019) in R (R Core Team 2021). This function colors the branches depending on the support value associated with that position. Hence, while all analyses presented in this paper included fossils as terminal taxa, RoguePlots do not show the fossil in question as a tip in the phylogeny.

## RESULTS

### Constrained Analyses

The phylogenetic placements of fossils obtained with parsimony, ML, and BI are summarized in Table 3. The results are generally consistent across all methods, resulting in similar positions for the fossils but with slight variations. The *CMP-1-Morph* and *CML-1-Morph* analyses, retrieved the same overall placement of fossils but differed slightly in their optimal position. For example, the most parsimonious (MP) position of *Canrightia resinifera* corresponds to the family Stemonaceae (Pandanales), whereas positions one step longer than the most parsimonious one (MP+1) are in Chloranthaceae (Chloranthales), and in Thismiaceae (Dioscoreales) (Fig. S7a, Fig. S7b). In turn, the best supported maximum likelihood position is restricted to Chloranthaceae (70-80 BS) (Fig. S7c). For other ML approaches, the results were similar. However, *CML-24-TE* (Fig. S7e) associated fossils to a larger number of families than *CML-1-Morph* (Fig. S7c) and *CML-24-Morph* (Fig. S7d), as can be observed for fossils such as *Archaestella verticillatus*, *Bertilanthus scanicus*, *Dakotanthus cordiformis*, and *Florissantia quilchenensis*, among others (Table 3).

**Table 3.** Summary of potential positions of fossils within major lineages and/or orders obtained across methods. * Indicates moderate confidence and ** high confidence

| Fossil | CML-1-Morph | CML-24-Morph | CML-24-MorphTE | UML-24-TE | UBI-24-TE |
| --- | --- | --- | --- | --- | --- |
| *Archaeanthus linnenbergeri* | ANA grade, Magnoliidae | ANA grade, Magnoliidae | ANA grade, Magnoliidae | ANA grade, Magnoliidae | Magnoliales* |
| *Archaeostella verticillatus* | Trochodendrales** | Trochodendrales* | ANA grade, Ranunculales, Trochodendrales | ANA grade, Ranunculales, Trochodendrales | Trochodendrales* |
| *Bertilanthus scanicus* | Saxifragales* | Saxifragales* | Saxifragales, Apiales, Caryophyllales | Saxifragales, Apiales, Asterales | Saxifragales* |
| *Canrightia resinifera* | Chloranthales, Monocots | ANA grade, Chloranthales, Magnoliidae, monocots | ANA grade, Chloranthales, Magnoliidae, monocots | ANA grade, Chloranthales, Magnoliidae, monocots | Piperles** |
| *Carpestella lacunata* | ANA grade, Magnoliidae | ANA grade, Magnoliidae | ANA grade, Magnoliidae | ANA grade, Magnoliidae | Amborellales** |
| *Cecilanthus polymerus* | ANA grade, Chloranthales, Magnoliidae | ANA grade, Chloranthales, Magnoliidae | ANA grade, Chloranthales, Magnoliidae | ANA grade, Chloranthales, Magnoliidae | ANA grade* |
| *Chloranthistemon endressii* | Chloranthales** | Chloranthales, Magnoliidae | Chloranthales* | Chloranthales** | Monocots |
| *Dakotanthus cordiformis* | Sapindales, Picramniales | Sapindales, Fabales, Crossosomatales | Sapindales, Fabales, Crossosomatales | Sapindales, Fabales, Malpighiales, Crossosomatales | Oxalidales, Fabales, Sapindales |
| *Divisestylus brevistamineus* | Saxifragales, Cornales | Saxifragales* | Saxifragales* | Saxifragales* | Saxifragales* |
| *Florissantia quilchenensis* | Malvales, Sapindales | Malvales, Malpighiales, Sapindales | Sapindales, Huerteales, Brassicales | Sapindales, Malpighiales, Huerteales, Malvales | Santalales, Malpighiales |
| *Kajanthus lusitanicus* | Ranunculales* | Ranunculales* | Ranunculales* | Ranunculales* | Ranunculales* |
| *Mabelia connatifila* | Malpighiales** | Malpighiales** | Malpighiales** | Malpighiales, monocots, Magnoliidae | Piperles* |
| *Mauldinia mirabilis* | Laurales* | Laurales* | Laurales* | Laurales* | Laurales* |
| *Microvictoria svitkoana* | ANA grade and Magnoliidae | ANA grade and Magnoliidae | ANA grade and Magnoliidae | ANA grade and Magnoliidae | ANA grade and Magnoliidae |
| *Paleoclusia chevalieri* | Sapindales,<br>Huerteales, Malvales | Sapindales, Malvales | Sapindales, Malvales | Sapindales,<br>Malvales,<br>Malpighiales | Ericales, Malpighiales,<br>Sapindales |
| *Paradinandra suecica* | Sapindales, Malvales | Brassicales, Malvales | Brassicales, Malvales | Sapindales,<br>Malvales, Ericales | Caryophyllales,<br>Sapindales |
| *Pentapetalum trifasciculandricus* | Dilleniales, Ericales | Dilleniales, Malpighiales,<br>Ericales | Dilleniales,<br>Caryophyllales,<br>Ericales | Dilleniales,<br>Malpighiales,<br>Ericales | Sapindales,<br>Caryophyllales |
| *Platydiscus peltatus* | Myrtales,<br>Crossosomatales | Myrtales, Sapindales,<br>Crossosomatales | Myrtales,<br>Saxifragales,<br>Sapindales | Cucurbitales,<br>Myrtales,<br>Sapindales,<br>Saxifragales | Myrtales* |
| *Quadriplatanus georgianus* | Proteales* | Proteales, Ranunculales,<br>ANA grade,<br>Ceratophyllales | Proteales,<br>Ranunculales, ANA<br>grade | Proteales,<br>Ranunculales, ANA<br>grade, Saxifragales | Malpighiales** |
| *Rariglanda jerseyensis* | Geraniales, Myrtales,<br>Sapindales | Geraniales, Sapindales,<br>Myrtales, Brassicales,<br>Malpighiales | Sapindales,<br>Geraniales,<br>Saxifragales,<br>Crossosomatales | Sapindales, Ericales | Malpighiales** |
| *Saururus tuckerae* | Piperales, Monocots | Piperales* | Piperales,<br>Chloranthales,<br>monocots | Piperales,<br>Chloranthales,<br>monocots | Santalales, monocots |
| *Spanomera mauldinensis* | Buxales, Gunnerales,<br>Trochodendrales,<br>Saxifragales | Buxales, Gunnerales, | Buxales, Gunnerales,<br>Ranunculales | Buxales, Gunnerales,<br>Ranunculales,<br>Saxifragales | Buxales* |
| *Tylerianthus crossmanensis* | Saxifragales,<br>Trochodendrales | Saxifragales, Myrtales,<br>Dilleniales, Asterales,<br>Lamiales | Saxifragales,<br>Lamiales, Solanales | Saxifragales,<br>Lamiales, Asterales | Saxifragales,<br>Escalloniales, Apiales |
| *Virginianthus calycanthoides* | Magnoliales,<br>Laurales | ANA grade and<br>Magnoliidae | ANA grade and<br>Magnoliidae | ANA grade and<br>Magnoliidae | Magnoliales* |

### Unconstrained Analyses

Phylogenetic placements of fossils derived from unconstrained ML and BI analyses using total evidence are summarized in Table 3. The unconstrained ML total evidence phylogeny for extant and fossil species is shown in Fig. 1. Including fossils in the ML and BI analyses using total evidence did not change the topology among extant species, as inferred by Ramírez-Barahona et al. (2020) using only molecular data. However, the use of a total evidence data set resulted in substantial decreases in the support values for most extant clades, as well as decreased resolution at deeper phylogenetic levels, presumably due to high uncertainty in the position of some fossils (Fig. S1 and Fig. S2). The placements of fossils were found to be weakly or moderately supported (indicated by blue and green triangles in Fig. 1), except for *Chloranthistemon endressii* (red triangle in Fig 1).

**Figure 1.**
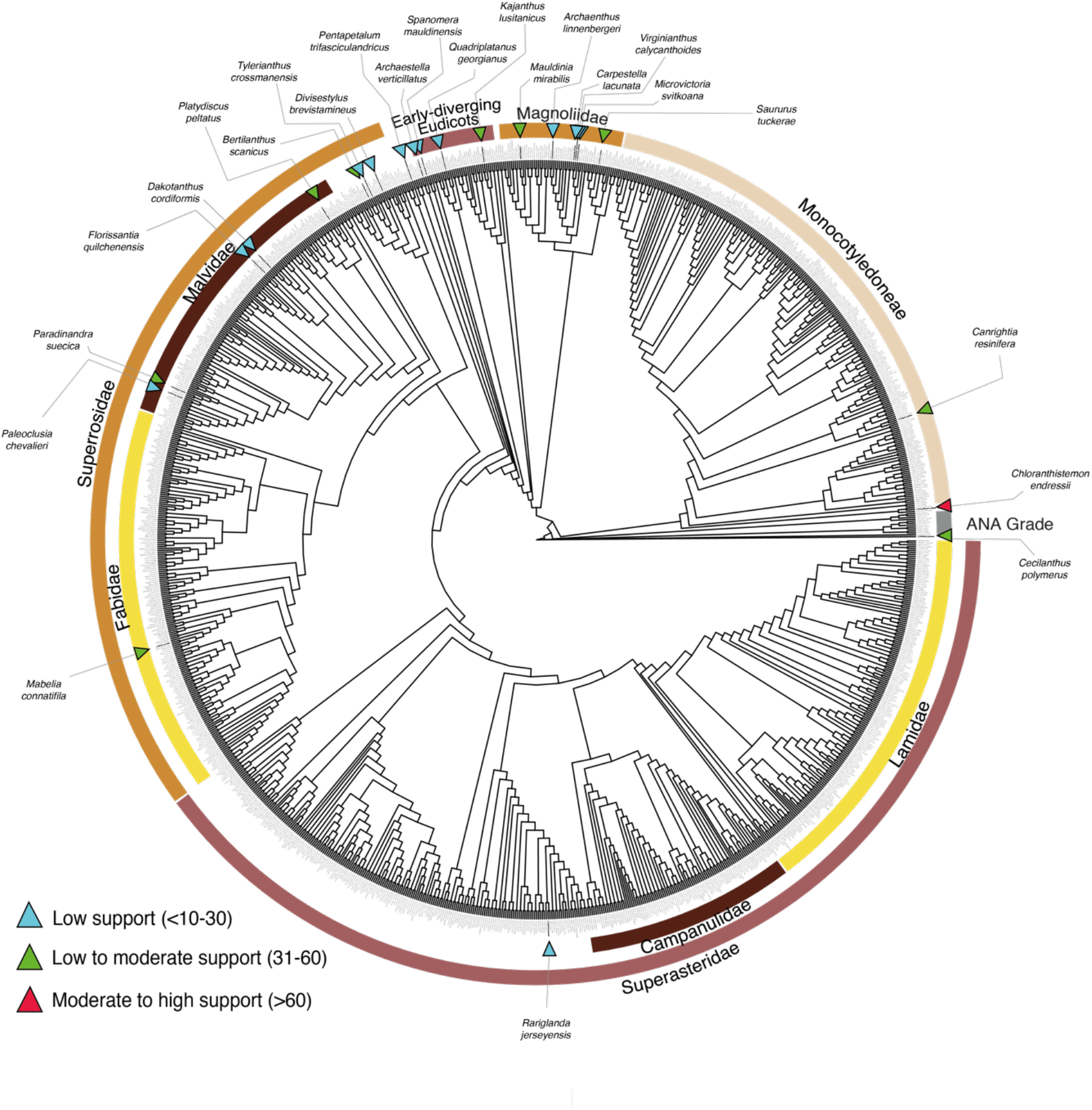
Maximum likelihood tree obtained with total-evidence data indicating the position of 1201 extant species and 24 fossil flowers included in this study. The support values were obtained from bootstrap replicates. Blue triangles indicate low support (<10-30%); green triangles indicate low to moderate support (31-60%); red triangles indicate moderate to high support values (<60%). Although this analysis produced a phylogram (Fig S1), it is presented here as a cladogram for the sake of clarity. For detailed results of each fossil, including alternative positions, see Supplementary Information.

Major phylogenetic inconsistencies were obtained in the unconstrained BI total evidence analysis with respect to parsimony and ML analyses (Table 3). *Chloranthistemon endressii,* for instance, was associated with Chloranthaceae in parsimony and ML analyses (*CML-1-Morph* (70-80 BS); *CML-24-Morph* (40-50 BS); *CML-24-TE* (60-70 BS); *UML-24-TE* (60-70 BS)); but in the BI analysis, this fossil attached to the branch of *Pogonia japonica* (Orchidaceae) (40-50% PP). These apparent anomalous positions were also observed for other fossils such as *Florissantia quilchenensis*, *Mabelia connatifila*, *Paradinandra suecica,* and *Tylerianthus scanicus*. Additionally, we observed that some fossils were linked with high support to branches corresponding to parasitic extant species. For example, *Quadriplatanus georgianus,* a fossil with distinctive attributes of Platanaceae (Proteales), is placed on the branch leading to *Rafflesia keithii* (70-80% PP) (Fig. S19f), belonging to the parasitic family Rafflesiaceae in the Malpighiales; while *Canrightia resinifera* with distinctive attributes of Chloranthaceae and Piperales, was placed with high support (80-90%PP) along the branch leading to *Hydnora africana* (Aristolochiaceae, Piperales), another parasitic plant (Fig. S4f).

### Uncertainty in the Placement of Fossils

Due to the high uncertainty associated with the placement of most fossils, we recognize the importance of considering all of the relationships inferred for fossils, rather than a single most supported position. We categorized the position of fossils into distinct levels of confidence, depending on the number of estimated distinct phylogenetic positions, and the associated support. We attributed “high confidence” to the placement of fossils attached to a single branch with high support (BS and PP) (e.g., *Chloranthistemon endressii* attached to the branch leading to *Sarcandra chloranthoides*, Fig. 2a). We attributed “moderate confidence” to the placement of fossils attached to a single branch in different analyses with moderate support (e.g., *Cecilanthus polymerus*, Fig. 2b), or attached to closely related branches (e.g., within the same family or order) with low support (e.g., *Kajanthus lusitanicus*, Fig. 2c; or *Bertilanthus scanicus*, Fig. 2d). Finally, we attributed “low confidence” to the placement of fossils that attached to several branches belonging to widely divergent clades with weak support (e.g., *Carpestella lacunata,* Fig. 4).

**Figure 2.**
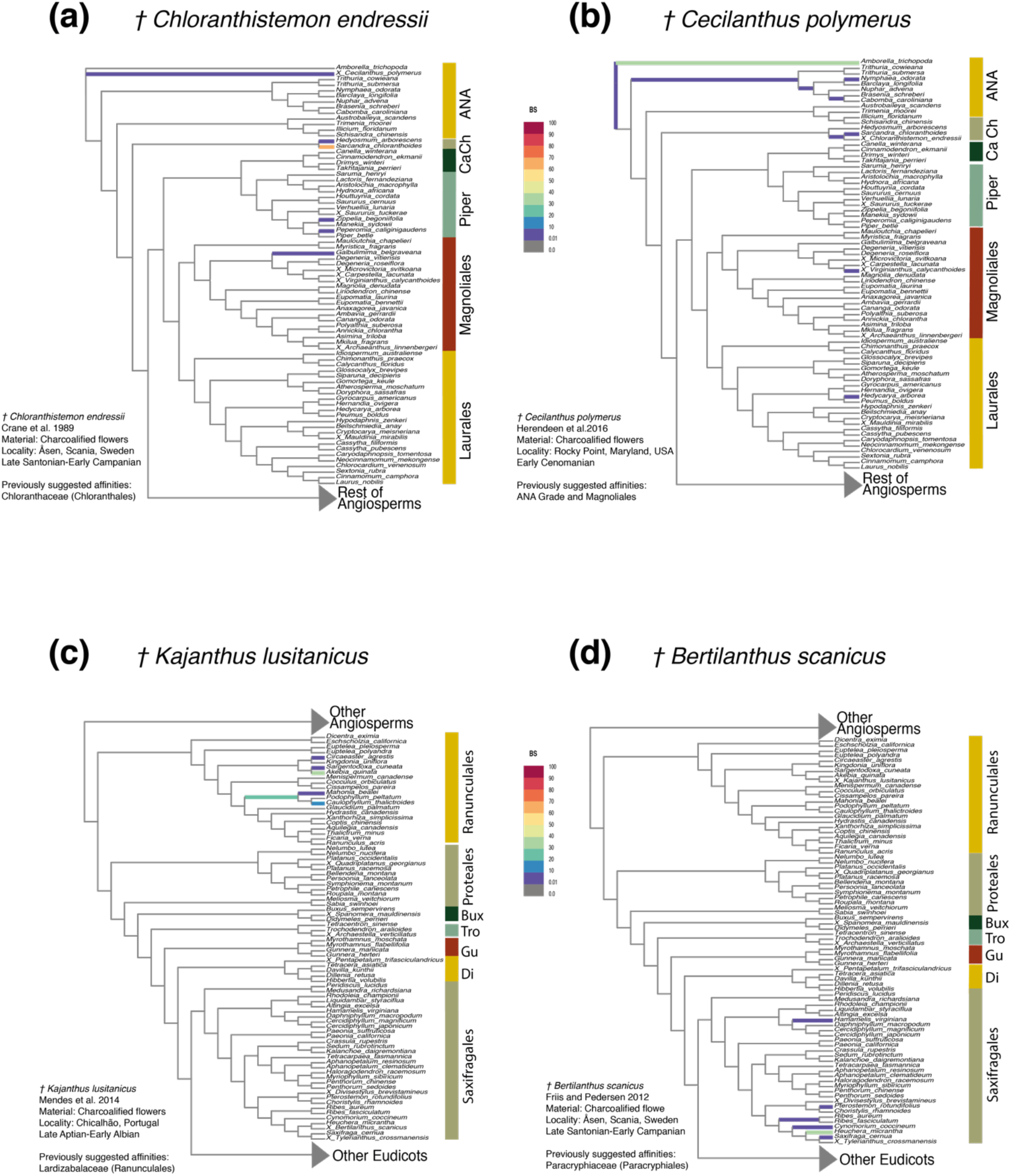
Partial RoguePlots derived from the total-evidence maximum likelihood analysis. The plots indicate all alternative phylogenetic positions of fossils and their associated uncertainty. The branches are colored according to the bootstrap values (BS values in color legend) associated with the attachment of the fossil to the branch. a) *Chloranthistemon endressii*; b) *Cecilanthus polymerus*; c) *Kajanthus lusitanicus*; d) *Bertilanthus scanicus*. Abbreviations: ANA, ANA grade (*Amborella*, Nymphaeales, Austrobaileyales); Ca, Canellales; Ch, Chloranthaceae; Piper, Piperales; Bux, Buxales; Tro, Trochodendrales, Gu, Gunnerales; Di, Dilleniales.

**Figure 3.**
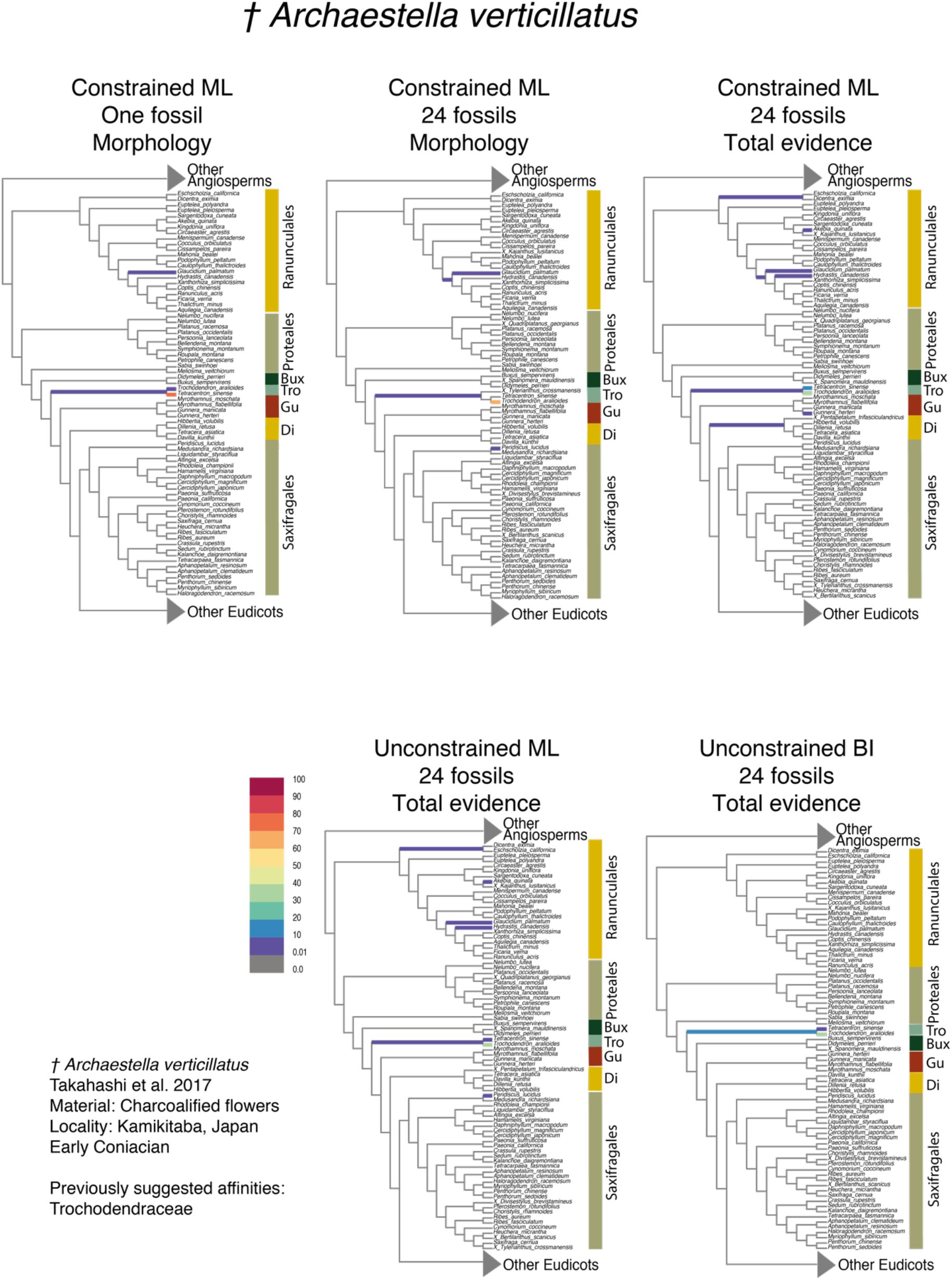
Partial RoguePlots indicating the phylogenetic placement of *Archaestella verticillatus* obtained from the five parametric methods implemented. The branches of the tree are colored according to the bootstrap values associated with the positions retrieved (BS values in color legend). Abbreviations: Bux, Buxales; Tro, Trochodendrales; Gu, Gunnerales; Di, Dilleniales.

**Figure 4.**
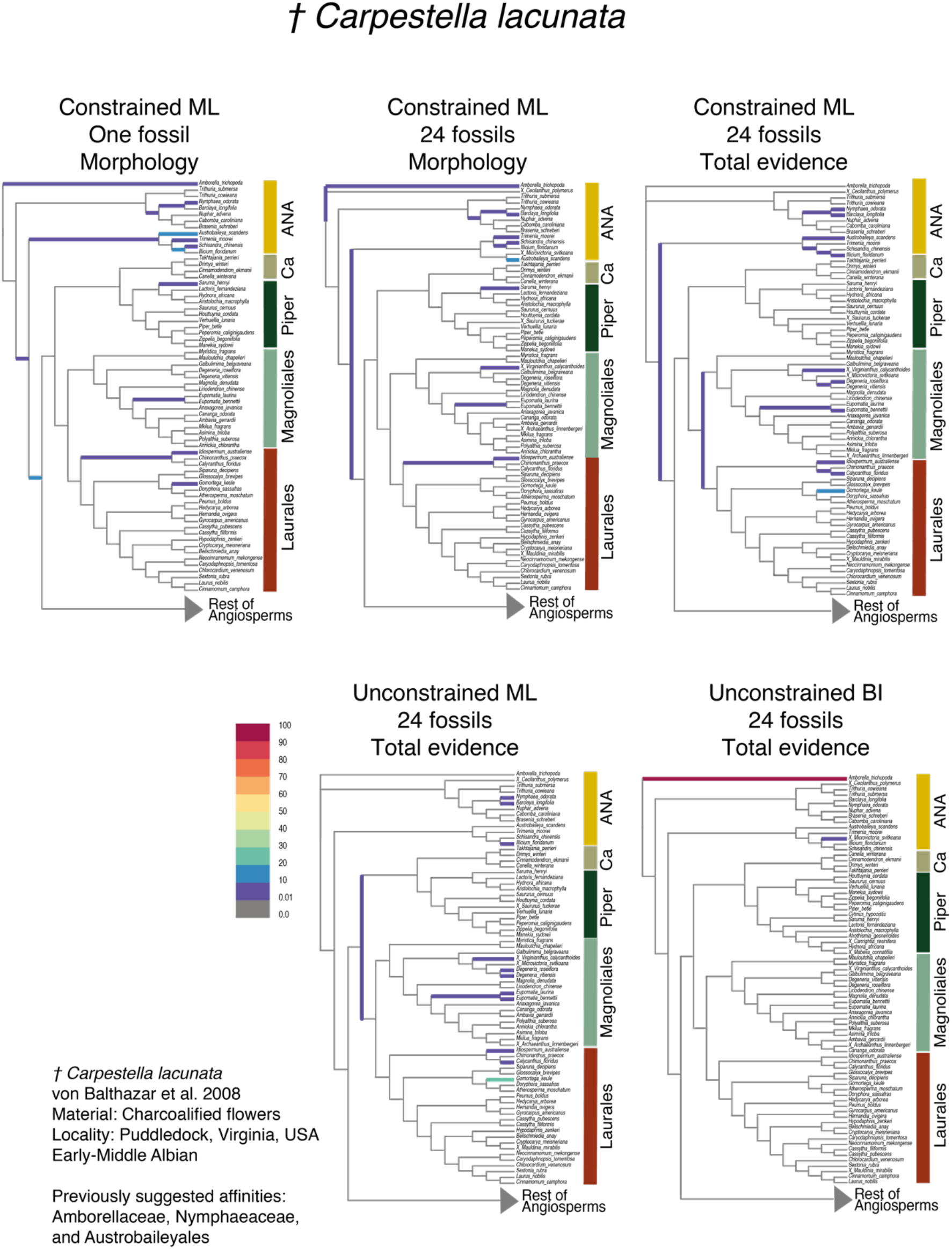
Partial RoguePlots indicating the placement of *Carpestella lacunata* obtained from the five parametric methods implemented. The branches of the tree are colored according to the bootstrap values associated with the positions retrieved (BS values in color legend). Abbreviations: ANA, ANA grade (*Amborella*, Nymphaeales, Austrobaileyales); Ca, Canellales; Piper, Piperales.

The phylogenetic placement of some fossils was recovered with similar levels of confidence across different phylogenetic analyses. For instance, *Archaestella verticillatus* was moderately supported as linked to the branch leading to *Trochodendron aralioides* (Trochodendraceae) in all analyses (Fig. 3). In contrast, the placement of some fossils exhibited substantial differences in support across different approaches. In general, BI analyses recovered placements that were restricted and strongly supported in particular positions. For example, *Carpestella lacunata* showed high phylogenetic uncertainty (low confidence) in ML analyses (Fig. 4), being attached with low support (<10-30 BS) to several independent branches across the ANA grade (Amborellaceae, Nymphaeaceae, and Austrobaileyaceae) and the orders Piperales, Magnoliales, and Laurales within the Magnoliidae clade. Conversely, in the total evidence BI analysis, the same fossil was attached to *Amborella trichopoda* with high support (90-100% PP). A similar pattern was observed in *Cecilanthus polymerus* (Fig. S6f).

## DISCUSSION

### Methods to Place Fossils into Phylogenies

In this study, we estimated the placement of fossil flowers across the angiosperm phylogeny using different optimization criteria and showed that, regardless of method and datasets, the different methods produced largely similar topological placements of fossils. In the past, the phylogenetic placement of angiosperm fossils has been mostly based on parsimony approaches focusing on individual fossils by using only morphological characters (e.g., Doyle and Endress, 2014; Eklund et al. 2004). Analyses of a total evidence matrix allow for the direct interaction between morphological and molecular data in estimating fossil phylogenetic relationships. Previously, Bayesian analyses of total evidence matrices have been reported to yield high accuracy in the topology in studies that incorporate extinct tips (Guillerme and Cooper 2016). In contrast to these findings, BI analyses here resulted in unexpected phylogenetic placements for some fossils, at odds with the more consistent results obtained from all other approaches tested in this study. Hence this method might not be appropriate under some conditions, such as in the incorporation of multiple extant terminals with high proportions of missing molecular data. Thus, based on our results, we recommend the implementation of total evidence analysis using maximum likelihood.

We advocate the use of broad-scale phylogenetic analyses as an important tool when estimating the placement of fossils. This approach may offer some key advantages in the assignment of fossils among living species. For example, this can reveal relationships that were not immediately obvious, and avoid potential confinement of fossils in potentially erroneous clades imposed *a priori*. As also stressed by Schönenberger et al. (2020), we consider that a more reliable estimation of the phylogenetic placement of any given fossil may be obtained in a two-step process. The first step relies on broad-scale analysis to identify different potential affinities of the fossil among major clades, and a second step may then focus on a particular clade and involve a more detailed morphological analysis and including a denser taxonomic and character sampling. In addition, we consider that the visualization of uncertainty associated with phylogenetic placement of fossils through RoguePlots (or other approaches), is critically important. This can help us to identify positions that are more strongly supported, and consider this information and associated uncertainty in subsequent analyses, for example during the implementation of taxonomic constraints of fossils in some dating approaches (i.e., fossilized birth-death process).

### Missing Data and Morphological Signal in the Integration of Fossils into Phylogenies

Total evidence analyses are often afflicted with different sources of missing data, which has been considered potentially problematic for achieving accurate phylogenetic estimations. Indeed, understanding the impact of missing data has been an important subject of active study (Kearney and Clark 2003; Wiens 2003; Magallón 2007; Manos et al. 2007; Wiens and Morrill 2011; Guillerme and Cooper 2016; Mongiardino Koch et al. 2021). Fossils represent a particularly challenging case, because they lack molecular information and the number of traits that can be scored is typically limited due to the fragmentary nature of preservation, especially in plants. However, studies of empirical and simulated data have demonstrated that including fossils can have a strong positive impact on phylogenetic estimation; fossils potentially allow to break long branches and may provide character combinations not present among extant species, which may produce significant topological rearrangements. However, a high proportion of missing data for fossils can also strongly decrease the accuracy and precision of the recovered topology, especially among deep nodes (Mongiardino Koch and Parry 2020; Mongiardino Koch et al. 2021).

Our results corroborate previous findings suggesting that the number of morphological characters scored for fossils does not have an impact on their estimated phylogenetic placement (Manos et al. 2007). For example, *Chloranthistemon endressii* is a highly incomplete fossil flower, of which only the three-lobed androecium was recovered while other parts of the flower, i.e., the perianth and the gynoecium, remain unknown (Crane et al. 1989). Given the set of floral characters we used, *C. endressii* inevitably has a high proportion of missing (or inapplicable) data; we were able to score only 12 out of 30 morphological characters. Despite this large amount of missing data, we found this fossil to be consistently placed in Chloranthaceae (with the exception of our *UBI-24-TE* analysis), in agreement with previous studies (Crane et al. 1989; Eklund et al. 2004; Schönenberger et al. 2020). Counterintuitively, the phylogenetic placement of some fossils for which many characters were scored (e.g., *Dakotanthus cordiformis*, *Florissantia quilchenensis*, *Rariglanda jerseyensis*) remained generally equivocal, being associated with multiple distantly related families across all analyses. This suggests that the effect of missing data on obtaining reliable positions is reduced for fossils displaying a particular set of highly distinctive morphological characters, especially if those characters are synapomorphies of particular clades (Manos et al. 2007). For example, the androecium of *C. endressii*, consisting of three stamens that are fused at the base, and with prominent connective extensions, represents a unique combination of characters shared only within modern Chloranthaceae (Crane et al. 1989).

An additional source of missing data is the often incomplete knowledge of the morphology of extant species. For instance, some studies make use of previously published morphological matrices, whose taxon sampling does not fully overlap with the molecular matrices, thereby resulting in a significant amount of missing data (e.g., Arcila et al. 2015). Based on simulations, Guillerme and Cooper (2016) proposed that the morphological data collected for extant species plays a major role in increasing the accuracy of phylogeny estimation under a total evidence approach. Our morphological matrix contained 32% of morphological characters missing, including inapplicable characters, but we did not detect any suspicious patterns of fossil attachment to extant species with high or low proportions of missing morphological data. Despite the considerable expansion of our taxonomic sampling of extant species (∼50%), compared to our previous study (Schönenberger et al. 2020), we obtained equivalent levels of uncertainty in the placement of fossils. This indicates that future increase in taxon sampling is unlikely to resolve the phylogenetic position of several fossil flowers.

Missing molecular data of extant species might also affect the position retrieved for fossils. The total evidence BI analysis reconstructed particular fossils as associated to parasitic lineages, which include species with large amounts of missing molecular data. The interaction between morphological and molecular data in the total evidence analyses, and the behavior of fossils attaching to terminals that are missing molecular data needs to be further investigated in future analyses using simulations. Even though we did not detect an overall negative impact of missing data on fossil placement, missing data (particularly molecular) may have a substantial effect on branch-length estimation, which is crucial for the accurate estimation of divergence time and ancestral characters in total evidence dating approaches.

### Phylogenetic Placement of Fossil Flowers within a Large Angiosperm Phylogeny

Our results show that the majority of fossil flowers were placed within orders or families that had been previously suggested based on detailed morphological comparisons (e.g., *Archaestella verticillatus* in Trochodendraceae, Takahashi et al. 2017; *Quadriplatanus georgianus* in Platanaceae, Magallón-Puebla et al. 1997; *Spanomera mauldinensis* in Buxaceae, Drinnan et al. 1991). Moreover, some fossil placements coincide with results from parsimony clade-specific analyses, such as the positions of *Divisestylus brevistamineus* in Saxifragales (Hermsen et al. 2003), *Saururus tuckerae* in Piperales (Smith and Stockey 2007), and *Chloranthistemon endressii* in Chloranthales (Eklund et al. 2004). These results suggest that a priori hypotheses about the placement of certain fossils based on expert observations are highly valuable, but also highlight the ability of our approach in recovering similar placements even with a relatively low number of floral traits. We also obtained fossil positions similar to previous broad-scale analyses based on parsimony, comparable to the *CMP-1-Morph* and *CML-1-Morph* approaches used here, such as the placement of *Carpestella lacunata* in Nymphaeaceae and Austrobaileyaceae, two families within the ANA grade (Fig. 4; Table 3; Fig. 3 in Doyle and Endress, 2014; Fig. 4 in von Balthazar et al. 2008).

On the other hand, for some fossils we obtained new and sometimes unexpected phylogenetic placements across all phylogenetic analyses. For instance, *Mabelia connatifila* was previously suggested to belong to Triuridaceae in the monocot order Pandanales, based on morphological comparison and parsimony analyses (Gandolfo et al. 2002). Our results from parsimony and ML approaches instead consistently placed this fossil in family Salicaceae, nested in the eudicot order Malpighiales, with high support (Table 3; Figures S12a, S12b, S12c, S12d, S12e, S12g). Another example is the consistent placement of *Bertilanthus scanicus* in Saxifragaceae (Saxifragales, Pentapetalae) across all approaches (Table 3), in contrast with suggested affinities to the distantly related Paracryphiaceae (Paracryphiales) (Friis and Pedersen, 2012).

The phylogenetic positions of some fossil flowers remain uncertain, specifically for those previously assigned to different subclades of Pentapetalae. Our morphological dataset mainly comprises characters of floral organization or groundplan (bauplan) (Endress 1994), such as the number of organs in whorls (merosity) and their arrangement. These characters are relatively stable at deeper phylogenetic levels among angiosperms clades and within Pentapetalae, especially because these structural attributes were apparently established at the onset of the diversification of eudicots, and remained mostly unchanged during subsequent evolution (Endress 1994; Sauquet et al. 2017). In particular, most members of Pentapetalae display flowers characterized by a whorled phyllotaxis, a differentiated perianth (calyx and corolla), and a pentamerous (or tetramerous) perianth and androecium (Endress 2010; Sauquet et al. 2017). Fossils such as *Platydiscus peltatus, Dakotanthus cordiformis, Paradinandra suecica,* and *Paleoclusia chevalieri* all share this generalized and conserved structure and are therefore less likely to be found in restricted unique phylogenetic positions in angiosperm-wide analyses. In order to obtain stronger signals at shallower phylogenetic levels, additional characters at the level of floral construction and mode are needed (e.g., size, shape, and conformation of complex and novel structures).

### Future Analyses and Perspectives

Our broad-scale phylogenetic analyses of fossil flowers, derived from a comprehensive and curated new morphological dataset for living species across all angiosperm families, represent a significant step forward for paleobotanical investigation. We also document the outcome of different strategies for combining fossil and extant species in phylogenetic analyses, which resulted in varying levels of uncertainty in fossil relationships. Our approach could be used in subsequent studies, for example, to estimate the placement of a larger number of fossil flowers, including newly discovered or undescribed fossils of unknown affinities, with respect to extant families and orders as recommended by Sauquet and Magallón (2018). This approach is also useful to corroborate previous assignments of fossils based on morphological assessments and to evaluate alternative placements across the phylogeny. Our morphological matrix and analytical approaches, especially those indicating the level of phylogenetic uncertainty associated to positions, will be useful to estimate the general phylogenetic relationships of a large number of previously described fossil flowers. The morphological data set used in our analyses is the result of considerable effort in scoring and curating data covering all currently expected angiosperm families. Further improvement and extensions to this data set in the future will allow for greater certainty about the phylogenetic placement of fossil flowers. Therefore, we suggest to conduct similar efforts on building and expanding morphological datasets, not only in terms of the number of living species that are scored, but most importantly in number and type of characters (e.g., incorporating detailed floral traits or vegetative traits).

Integration of fossils into extant phylogenies has become more popular since the implementation of the Fossilized Birth-Death model (Heath et al. 2014) and dating methods that can incorporate total-evidence (Ronquist et al. 2012, 2016; Zhang et al. 2016; Gavryushkina et al. 2017). The latter approach integrates molecular, morphological, and stratigraphic data of fossils into a single framework with extant species to simultaneously infer phylogenetic relationships and divergence times. Recently, King (2021) revealed that stratigraphic data and morphological data are reciprocally informative in the estimation of phylogenetic relationships, hence, we consider it desirable to evaluate the effect of ages in the placement of fossil flowers by conducting tip-dating analyses. Additionally, we recommend to consider the uncertainty in the placement of fossils before selecting those fossils that can be incorporated in macroevolutionary inferences, such as node calibrations in molecular clock analyses (as shown by Klopfstein and Spasojevic 2019), and to reduce the possibility of calibrating an internal node with a fossil with a phylogenetically incorrect or uncertain position.

Phylogenetic approaches that integrate fossils and extant species are essential to provide a more comprehensive picture of deep time macroevolutionary processes. Examples of such studies include the investigation of fruit evolution through time in Fagales (Larson-Johnson 2016), understanding the timing and diversification dynamics of Cetacea (Lloyd and Slater 2021), or inferring the biogeographical history of Juglandaceae (Zhang et al. 2021) and *Hypericum* (Meseguer et al. 2015). The fossil record of angiosperms is diverse and relatively abundant since their initial appearance in the Early Cretaceous (Friis et al. 2010). As in any other group of organisms, the angiosperm fossil record is also biased in different ways (i.e., temporally, spatially, and phylogenetically, as discussed by Xing et al. 2016). However, we expect that the continued compilation of large-scale morphological data sets and the subsequent integration of fossil flowers (and other types of fossils) into the angiosperm phylogeny will lead to the elucidation of deep time patterns of floral evolution.

## SUPPLEMENTARY MATERIAL

Data available from the Dryad Digital Repository: http://dx.doi.org/10.5061/dryad.[NNNN]

## ACKNOWLEDGEMENTS

We are grateful to the University of Vienna for funding the eFLOWER server hosting the PROTEUS database. We thank the numerous colleagues who have contributed to building this new eFLOWER dataset over the past five years, in particular Elisabeth Reyes, Luka Kovac, and all the participants from the Oak Spring eFLOWER Summer School. Peter Crane, the Oak Spring Garden Foundation, and the Society of Systematic Biologists are gratefully acknowledged for their support of this Summer School. A.L.M thanks the Posgrado en Ciencias Biológicas, Universidad Nacional Autónoma de México, and the Consejo Nacional de Ciencia y Tecnología (CONACyT) for granting scholarship 708068. We thank Natalia Ivalú Cacho for comments on earlier drafts; Alfonso Delgado, Luis E. Eguiarte, Rebeca Hernández, and Adriana Benítez for feedback and helpful discussions during different stages of this project.

## AUTHOR CONTRIBUTIONS

A.L.M., H.S., J.S., M.v.B., and S.M. designed the study. A.L.M., J.S., M.v.B., C.G.M., and H.S. scored and curated the morphological data of extant and fossil taxa. S.R.B. provided the backbone phylogeny and the molecular alignment. A.L.M conducted the analyses and wrote this paper, with subsequent input from all co-authors.

